# Tuning the Structural Properties of a Single-Domain Antibody Scaffold for Improved Fibroblast Activation Protein Targeting

**DOI:** 10.64898/2026.03.11.711127

**Authors:** Kendahl L. Ott, Joseph P. Gallant, Ohyun Kwon, Adedamola Adeniyi, Kendall E. Barrett, Zachary T. Rosenkrans, Jason C. Mixdorf, Eduardo Aluicio-Sarduy, Jonathan W. Engle, Reinier Hernandez, Bryan P. Bednarz, Aaron M. LeBeau

**Affiliations:** Molecular and Cellular Pharmacology Program, University of Wisconsin School of Medicine and Public Health, Madison, Wisconsin 53705, United States; Department of Pathology and Laboratory Medicine, University of Wisconsin School of Medicine and Public Health, Madison, Wisconsin 53705, United States; Department of Medical Physics, University of Wisconsin School of Medicine and Public Health, Madison, Wisconsin 53705, United States; University of Wisconsin Carbone Cancer Center, University of Wisconsin School of Medicine and Public Health, Madison, Wisconsin 53792, United States; Department of Radiology, University of Wisconsin School of Medicine and Public Health, Madison, Wisconsin 53792, United States

**Keywords:** fibroblast activation protein (FAP), single-domain antibodies, nuclear imaging, cancer-associated fibroblasts (CAFs), theranostic, protein engineering

## Abstract

Fibroblast activation protein (FAP) is an attractive target for the development of cancer theranostics due to its selective expression on cancer-associated fibroblasts (CAFs). While a number of small-molecule FAP inhibitors (FAPIs) have been developed, few biologics have been investigated as FAP targeting vectors. Camelid-derived single-domain antibodies, or variable-heavy-heavy domains (VHHs), offer a compelling alternative, combining high affinity with versatile engineering options. In this study, we first identified a novel anti-FAP VHH, F7, from an affinity-matured camelid phage display library. To investigate how valency and molecular weight affected target engagement and *in vivo* properties, F7 was engineered into three formats: a monomer (F7), a tethered dimer (F7D), and an Fc-fusion protein (F7-Fc). All three were specific for FAP with the two bivalent constructs demonstrating picomolar affinity. Positron emission tomography imaging in FAP-positive xenograft models revealed distinct pharmacokinetic profiles across constructs with notable differences in tumor uptake and clearance. F7 had rapid uptake and clearance resulting in significantly higher tumor uptake than FAPI-46. Low molecular weight bivalent F7D demonstrated similar kinetics but was retained by the tumor resulting in a high tumor-to-blood ratio with secondary uptake limited to clearance organs. The largest construct, F7-Fc, resulted in the highest tumor uptake and allowed for longitudinal imaging. Absorbed dose calculations confirmed that tumors received significantly higher radiation doses compared to normal tissues. These findings demonstrate that tuning VHH scaffold size and valency can improve biodistribution and retention, establishing F7-based constructs as promising targeting vectors for FAP.

## 1. INTRODUCTION

The serine protease fibroblast activation protein (FAP) is a cell-surface marker of cancer associate fibroblasts (CAFs), a stromal cell type that is a critical component of the tumor microenvironment (TME).^1,2^ Under normal physiological conditions, the expression of FAP is limited to embryogenesis and at sites of wound healing and fibrosis.^1,3,4^ Normal fibroblasts do not express FAP; it is only after a disease state transition has occurred that CAFs will uniformly express FAP.^5^ The expression of FAP in the TME has been reported in almost every type of solid cancer.^6^ The extensive expression of FAP in the TME of solid tumors suggests it participates in a complex signaling network that may aid in cancer progression and metastasis. FAP expression has been shown to correlate with increased cancer motility and decreased survival rates in several cancer types.^7,8^ CAFs potentially driven by FAP contribute either directly or indirectly to the epithelial-mesenchymal transition (EMT) process at the tumor site. Importantly, FAP is known to promote an immunosuppressive TME in tumors.^1,9,10^ Recent studies have shown that the selective elimination of FAP-expressing CAFs leads to the secretion of proinflammatory cytokines and the increased infiltration of CD8+ T cells into the TME.^11,12^ For these reasons, FAP is regarded as an alluring target for the development of targeted theranostics to see and treat cancer.

As a member of the S9 prolyl oligopeptidase family, FAP cleaves the peptide bond after proline residues in peptides and proteins.^13,14^ While most members of the S9 family are either amino- or carboxypeptidases, FAP possess an extended substrate specificity pocket which allows it to act as both an endo- and exopeptidase.^15–17^ This extended substrate specificity has been leveraged in the development of numerous small-molecule FAP inhibitors (FAPIs) that demonstrate greater specificity for FAP compared to other members of the S9 family.^18,19^ Due to their size, FAPIs make a limited number of contacts with FAP and tend to have low nanomolar inhibition constants.^18,20^ Coupled with the low density of FAP on the surface of CAFs, it is likely that it will be difficult to make effective FAPI-based radiotherapies given their quick wash out and poor retention in tumors.^21–23^ Though they are more expensive to produce for clinical applications, antibodies represent an alternative targeting strategy for FAP. In particular, the binding domains of camelid antibodies (also known as nanobodies or variable-heavy-heavy domains (VHHs)) are promising candidates for theranostic development.^24^ At only 15kDa in size, VHHs are the smallest antigen binding domain derived from heavy-chain antibodies.^25^ These domains only have three complementarity-determining regions (CDRs), however, the CDR3 is often longer than that of a conventional antibody resulting in a convex paratope that can access recessed areas of a target.^25,26^ Additionally, they can be engineered into monovalent, bivalent, and multivalent formats because of their size and ease of expression.^27^

Here, we identified a novel anti-FAP VHH domain, termed F7, from an affinity-maturated VHH phage display library. We then performed a PET imaging study using F7 to determine how valency and molecular weight affect FAP engagement, pharmacokinetics, and localization to tumors *in vivo*. As a single VHH, F7 demonstrated rapid accumulation to the tumor resulting in greater uptake than a clinically relevant FAPI. By tethering two copies of F7 together, we created a high-affinity homodimer that quickly localized to the tumor and was retained with modest uptake. A high molecular weight Fc fusion protein of F7 resulted in a construct with the highest tumor uptake and retention. Our findings suggest that F7 is a potent targeting domain for FAP that could be exploited for future theranostic applications.

## 2. MATERIAL AND METHODS

Methods for VHH identification and characterization (phage display biopanning, ELISA, protein purification, biolayer interferometry, and flow cytometry) can be found in the Supporting Information section.

### 2.1. Animal Studies

Animal studies were performed under guidelines approved by the University of Wisconsin Research Animal Resources Center. All mice in the study were athymic nude-Foxn1nu (Envigo). For subcutaneous xenografts, 1×10^6^ cells were injected in a 1:1 PBS and Matrigel (Corning) mixture and injected into either the area above the hind leg or above the animal’s shoulder. Animals were split evenly between the FAP positive group receiving CWR-R1^FAP^ cells or the negative group receiving the parental CWR-R1 cells. Tumors were allowed to grow to at least 300mm^3^ before initiation of the imaging studies.

### 2.2. PET/CT Imaging

For nuclear imaging studies, either ^64^Cu or ^68^Ga was used, and chelator addition and radiolabeling followed established protocols. ^64^Cu was supplied by the University of Wisconsin Medical Physics Department (Madison, WI). VHH constructs were first modified with p-SCN-Bn-NOTA (NOTA, Macrocyclics), after which ^64^Cu was conjugated in a sodium acetate (NaOAc) buffered reaction (pH 5.0). The conjugation ratios were 25 μg VHH, 50 μg dimer, and 100 μg Fc fusion per 3.7 MBq of ^64^Cu. Constructs were incubated with buffered ^64^Cu at 37 °C with gentle agitation. To complete conjugation, Tween-20 was added to a final concentration of 0.005%, and the labeled constructs were purified using a PD-10 size-exclusion column pre-equilibrated with PBS. All microPET/CT imaging was performed on an Inveon μPET/CT scanner (Siemens Medical Solutions). For comparison of camelid VHH and small-molecule tracers, mice (n = 3 per experimental and control group) were injected via tail vein with ^68^Ga-FAPI-46 (average dose: 5.9 MBq). One-hour post-injection, mice were anesthetized with 2.5% isoflurane and PET/CT images were acquired. The same mice were then injected the following day with ^64^Cu-labeled F7 VHH (average dose: 5.7 MBq), followed by a 1-hour dynamic scan and a second acquisition at 4 hours post-injection. PET list-mode data were collected for 80 million counts using a gamma-ray energy window of 350–650 keV and a coincidence timing window of 3.438 ns. CT-based attenuation correction was performed over approximately 10 minutes (80 kVp, 1 mA, 220° rotation, 120 steps, 250 ms exposure) and reconstructed using a Shepp–Logan filter with 210 μm isotropic voxels. PET images were reconstructed using 3D ordered-subset expectation maximization (OSEM3DMAP; 2 iterations, 16 subsets) with a maximum a posteriori probability algorithm. Two-dimensional images and maximum intensity projections (MIPs) were generated in Inveon Research Workplace, and regions of interest (ROIs) were manually drawn and quantified using Inveon Research Workspace. Imaging studies for the F7 dimer and F7-Fc followed the same protocol, with adjusted acquisition times. For the F7 dimer, scans were acquired at 1, 4, and 24 h post-injection of ^64^Cu-labeled F7 dimer (average dose: 8.7 MBq). For F7-Fc, scans were acquired at 4, 24, and 48 h post-injection of ^64^Cu-labeled F7-Fc (average dose: 9.9 MBq). Image quantification was performed as described above. All graphs and statistical analyses were generated using GraphPad Prism.

### 2.3. Biodistribution

Acute *in vivo* biodistribution studies were conducted to evaluate the uptake of [^64^Cu]Cu-VHH constructs in mice bearing subcutaneous CWR-R1 or CWR-R1^FAP^ tumors. Mice (n=3 mice for each tissue per timepoint) were euthanized at each final imaging timepoint postinjection. Organs (including the tumor) were excised and counted on an automatic gamma-counter (Hidex). The total number of counts [counts per minute (cpm)] of each organ was compared with a standard syringe of known activity and mass. Count data were background- and decay-corrected, and the percentage injected dose per gram (%ID/g) for each tissue sample was calculated by normalization to the total amount of activity injected into each mouse. All graphs and statistical analyses were generated using GraphPad Prism.

### 2.4. RAPID *in vivo* dosimetry

Mice bearing CWR-R1^FAP^ xenografts (n=3) were administered [^89^Zr]Zr-F7-Fc. Longitudinal μPET/CT imaging (Inveon; Siemens) was performed at 4, 24, 48, 72, 96, 120, and 144 h post-injection. ROIs for the tumor, heart, liver, kidneys, and bone were manually delineated at each timepoint to generate time–activity curves (%ID/g). Voxel-wise absorbed dose rate estimates (Gy/s/MBq) were calculated using the Geant4-based Monte Carlo dosimetry platform, RAPID^38^, with subject-specific PET images as the source distribution and corresponding CT images for geometry and tissue density.^28^ The total mean absorbed dose (Gy/MBq) for each ROI was determined by integrating the absorbed dose rates using the trapezoidal method, assuming physical decay of ^89^Zr beyond the final imaging timepoint.

### 2.5. iQID

Mice (n=1 per timepoint) bearing CWR-R1^FAP^ xenografts were injected with [^89^Zr]Zr-F7-Fc and sacrificed at 24 h and 144 h post-injection. Tumors were excised, frozen in OCT, and sectioned at 15 µm using a Leica CM1950 cryostat. Sections were mounted on charged glass slides (Fisherbrand Superfrost Plus®) for iQID imaging. Three serial sections were prepared for the 24 h timepoint and two for the 144 h timepoint. Slides were scanned individually against the iQID detector window, separated by a mylar sheet (Ludlum Measurements 01-5859) and a beta-sensitive intensifying screen (Carestream BioMax TranScreen LE). Scanning was performed once per slide (7 total): 40 min for 24 h samples and 1 h for 144 h samples. After autoradiographic scanning, tissue sections were imaged and stitched using Infinity ANALYZE 7 with an Infinity2-5C CCD camera on an Olympus CKX41 microscope (2× objective). Histological, iQID, and shadowgraph images were acquired; shadowgraphs were generated by illuminating tissue samples with a phosphorescent disk inside the iQID enclosure. All images were imported into ImageJ, where SIFT or MOPS algorithms identified shared features between shadowgraphs and histology for rigid alignment. bUnwarpJ was then applied for deformable registration. Final registered histology images were generated in Adobe Photoshop and overlaid onto corresponding iQID autoradiographs in ImageJ to produce comprehensive iQID/histology overlays.

## 3. RESULTS

To identify an anti-FAP VHH for PET imaging, we used our naïve VHH phage display library that was developed in-house (diversity of 7.5 x 10^10^).^29,30^ After four rounds of screening against recombinant human FAP on magnetic beads, we identified 29 clones that bound FAP by ELISA. Of these clones, seven unique sequences were identified (Figure S1A, Table S1). All seven of the clones were specific for FAP by ELISA and did not cross-react with DPPIV (Figure S1B). DPPIV is a member of the S9 prolyl oligopeptidase family that shares 52% amino acid sequence homology with FAP.^23^ Based on ELISA data, a lead VHH was selected. The lead VHH, termed NbC, was expressed and purified, and its binding affinity was determined to be 70nM by biolayer interferometry (BLI) (Table 1). Affinity maturation was then performed on the VHH by site-specific mutagenesis at one amino acid in CDR1 and four amino acids in CDR3 (Figure 1A). Two additional rounds of biopanning against FAP were performed using the affinity matured sub-library with a diversity of 1.4 x 10^9^. Screening 384 clones from the first round yielded 93 strong binders (24.2%), while a subsequent screen of 480 clones from the second round identified 27 strong binders (5.62%) using a stringent ELISA (Figure S2A,B). Out of the 27 highest binding clones, 24 were revealed to be unique, largely in their CDR3 (Figure 1B). The top five VHH candidates were subsequently evaluated using a soluble dilution ELISA to assess their binding affinities. (Figure 1C, Table S2). Clone F7 was selected as our lead for further investigation based on its affinity for FAP (K_D_ – 9nM) and ease of expression in bacteria (>30mg/L yield).

**Figure 1.**
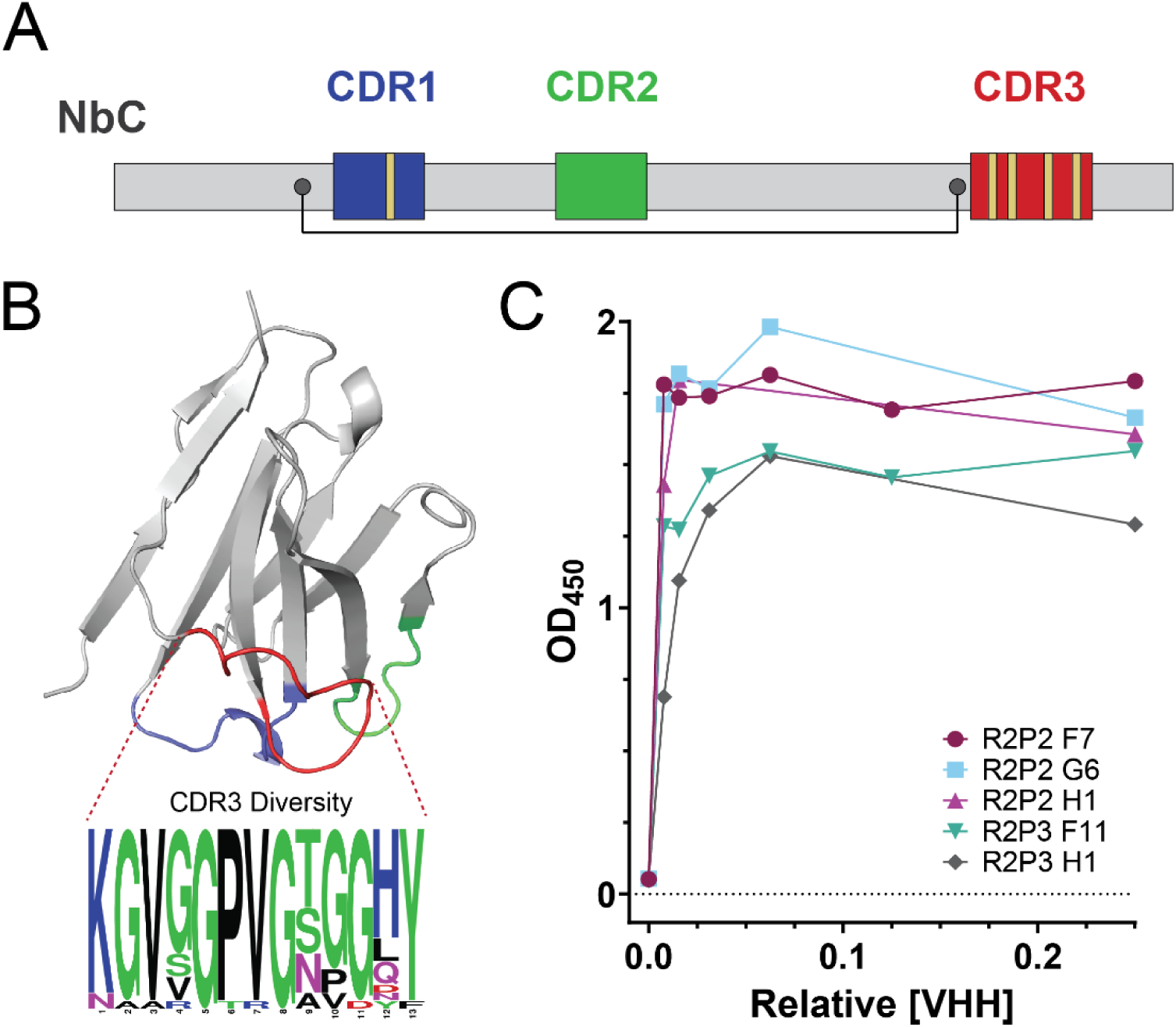
Identification of FAP-targeted VHHs domains from an affinity-maturated phage-display library. (A) Schematic of the VHH domain architecture with mutation sites annotated for the generation of the affinity-maturated phage-display library. Depicted are the framework regions (gray), CDR1 (blue), CDR2 (green), CDR3 (red), sites of targeted mutagenesis for the designated CDRs (yellow), and canonical cystine residues (dark gray). (B) Ribbon diagram of a representative VHH, colored as in (A) with CDR3 diversity illustrated by a sequence logo derived from top affinity-matured biopanning hits. (C) Binding curves of representative VHH clones measured by ELISA, measuring OD_450_ as a function of relative VHH concentration.

**Table 1.** Collated association rate (K_a_), dissociation rate (K_d_), and equilibrium dissociation constants (K_D_) of the indicated constructs for human FAP as determined by biolayer interferometry.

| Construct | $K_D$ | $K_a$ | $K_d$ |
| --- | --- | --- | --- |
| NbC | $7.08\text{E-}08 \pm 3.13\text{E-}09$ | $5.36\text{E}5 \pm 2.15\text{E}4$ | $3.82\text{E-}2 \pm 7.10\text{E-}4$ |
| F7 | $9.11\text{E-}09 \pm 1.37\text{E-}10$ | $8.90\text{E}5 \pm 1.28\text{E}4$ | $8.11\text{E-}3 \pm 3.59\text{E-}5$ |
| F7D | $2.90\text{E-}11 \pm 6.80\text{E-}12$ | $9.52\text{E}5 \pm 4.52\text{E}3$ | $2.77\text{E-}5 \pm 6.40\text{E-}6$ |
| F7-Fc | $3.87\text{E-}11 \pm 1.00\text{E-}12$ | $6.58\text{E}6 \pm 4.03\text{E}4$ | $2.55\text{E-}4 \pm 4.50\text{E-}6$ |

Data is shown as M with SD.

To investigate the effects of valency and molecular weight on engaging FAP and subsequent *in vivo* properties, two additional constructs using F7 were expressed and purified. F7 was engineered into a low-molecular weight bivalent homodimer (F7D) using a (G_4_S)_5_ linker to tether two VHHs together. We then grafted F7 onto a human Fc domain to create a high-molecular weight (80kDa) bivalent VHH-Fc fusion protein (F7-Fc). The K_D_ values for the bivalent constructs were found to be relatively similar at 29pM for F7D and 38pM for F7-Fc as measured by BLI (Figure 2A, Table 1). Most notably the two bivalent constructs had much slower rates of dissociation (K_d_) compared to the F7 monomer. No cross-reactivity was observed when the three constructs were screened by BLI against mouse FAP. Flow cytometry further highlighted the difference in affinity and K_d_ between the monovalent and bivalent constructs. All F7-engineered constructs showed specific binding to the FAP-positive prostate cancer cell line (CWR-R1^FAP^), whereas binding to the parental cell line was minimal. Notably, the two bivalent constructs (F7D and F7-Fc) showed greater binding to the CWR-R1^FAP^ cells across a series of concentrations (Figure 2B,C).

**Figure 2.**
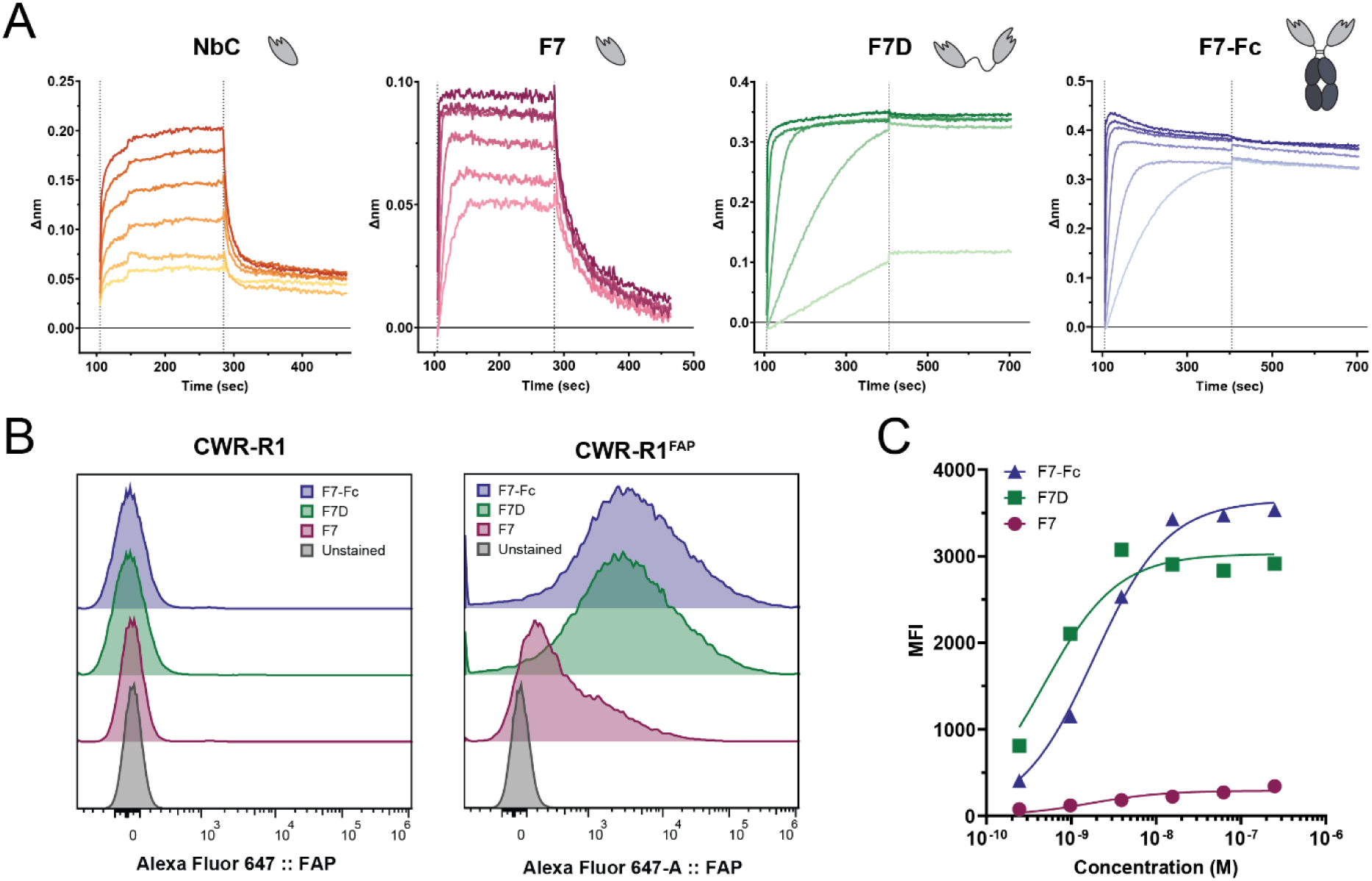
*In vitro* characterization of the lead anti-FAP VHH constructs. (A) BLI sensorgrams show sensors loaded with FAP and exposed to serial dilutions of the F7-based constructs, followed by dissociation in assay buffer. Representative VHH construct structures are displayed alongside their corresponding BLI traces. (B) Flow cytometry analysis of F7-based constructs labeled with Alexa Flour 647 to assess binding in FAP-null (CWR-R1) and FAP-positive (CWR-R1^FAP^) cell lines at 250 nM. Samples were compared to unstained cell controls. (C) Dose response curves for CWR-R1^FAP^ cells stained with varying concentrations of the F7-based constructs as assessed by flow cytometry.

After determining the *in vivo* specificity of [^64^Cu]Cu-F7 in FAP-positive CWR-R1^FAP^ and FAP-null CWR-R1 xenografts by PET and *ex vivo* biodistribution (Figure S3), we next compared the ability of our monovalent F7 to FAPI-46 at detecting FAP expression *in vivo*. Mice bearing FAP-positive CWR-R1^FAP^ xenografts were administered either [^68^Ga]Ga-FAPI-46 or [^64^Cu]Cu-F7. As anticipated for a small molecule and a protein with a molecular weight of 15kDa, the uptake for both imaging agents was rapid with image acquisition possible for both at the 1 h post-injection timepoint (Figure 3A). [^68^Ga]Ga-FAPI-46 showed rapid uptake in the FAP-positive xenograft with a %ID/g of 0.58% ± 0.143 at 1 h as determined by ROI analysis. Our low-molecular weight biologic [^64^Cu]Cu-F7 had tumor uptake two-fold higher than FAPI-46 in the same xenograft model with an average uptake of 1.1% ± 0.173 in at 1 h. [^64^Cu]Cu-F7 was also imaged and analyzed at 4 h with an average tumor uptake of 0.923% ± 0.125. The blood clearance of [^68^Ga]Ga-FAPI-46 was rapid with an *in vivo* ROI of only 0.104% ± 0.016. Similarly, [^64^Cu]Cu-F7 VHH had a blood uptake ROI of 0.357% ± 0.085 at 1 h and 0.196% ± 0.103 at 4 h. [^64^Cu]Cu-F7 had an expectedly high signal in the kidneys, 39.367% at 1 h and 29.567% at 4 h, consistent with clearance (Figure 3B).

**Figure 3.**
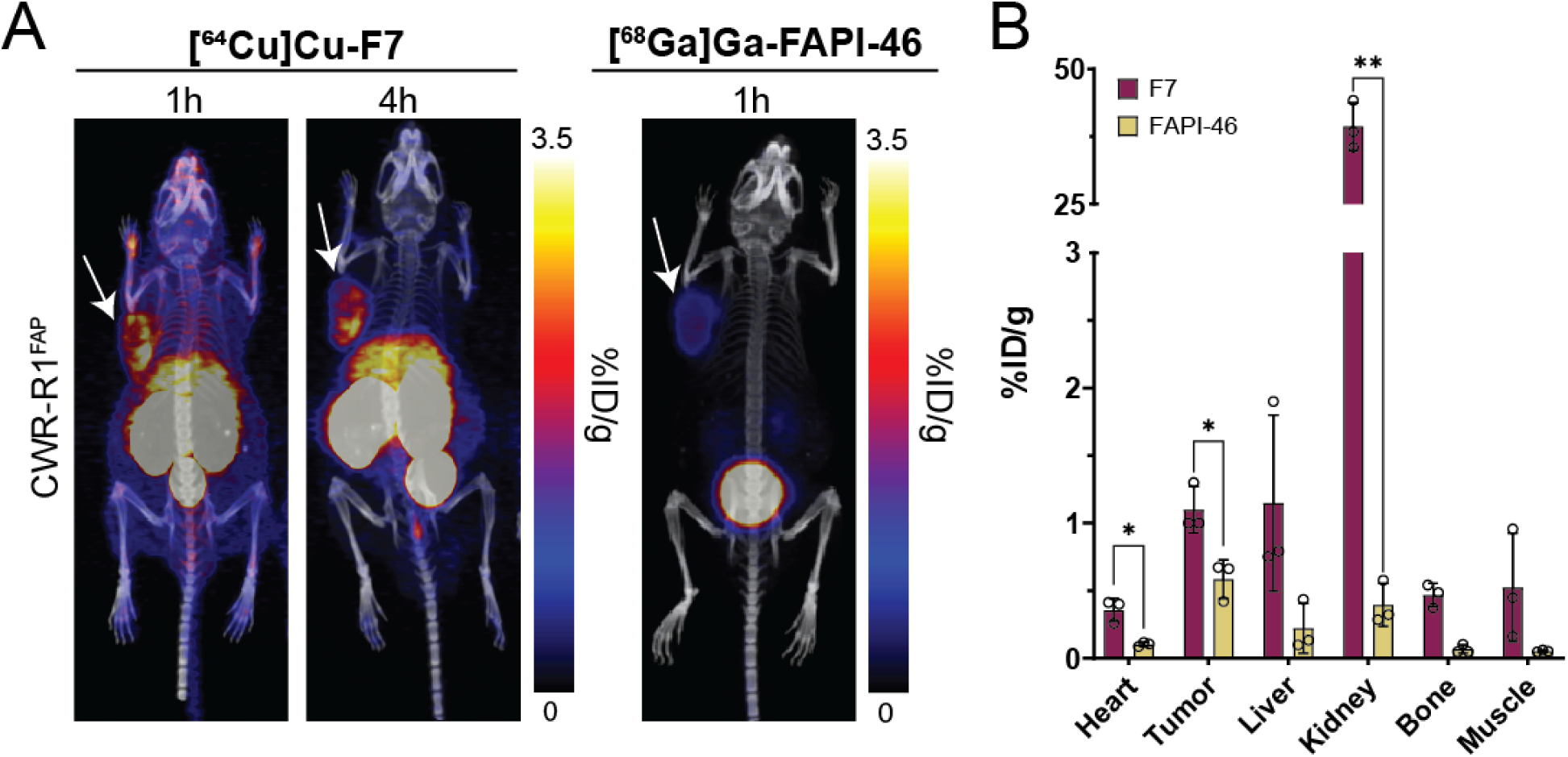
Direct comparison of F7 and FAPI-46 for nuclear imaging. (A) MIPs from PET/CT scans using [^64^Cu]Cu-F7 and the small molecule [^68^Ga]Ga-FAPI-46 in CWR-R1^FAP^ xenografts at 1 h and 4 h post-injection. Tumor indicated by white arrow. (B) Biodistribution at 1 h in mice bearing CWR-R1^FAP^ xenografts for F7 and FAPI-46. *P values*, * *P* ≤ 0.05, ** *P* ≤ 0.01 using unpaired T test with Welch correction.

The higher molecular weight F7 constructs, F7D and F7-Fc, were next investigated for their ability to image FAP in xenografts by PET. For this study, we decided to initially use copper-64 as our imaging isotope to be consistent among the different constructs tested. Mice bearing CWR-R1^FAP^ and CWR-R1 xenografts were injected with [^64^Cu]Cu-F7D or [^64^Cu]Cu-F7-Fc. [^64^Cu]Cu-F7D was imaged at 1 h, 4 h, and 24 h post-injection given its lower molecular weight of 28kDa (Figure 4A). Tumor uptake was rapid with the [^64^Cu]Cu-F7D in FAP-positive tumors having average ROI analysis values of 1.467%± 0.240, 1.433% ± 0.176, and 1.267% ± 0.133 at 1, 4, and 24 h respectively. Antithetically, in the FAP-null xenografts, tumor uptake was 0.581% ± 0.079 and 0.473% ± 0.041, 0.296% ± 0.030 again at 1, 4, and 24 h post-injection (Figure 4B). Tumor uptake increased with each later scan as [^64^Cu]Cu-F7D was retained in the FAP-positive tumors and cleared from the FAP-null tumors. The blood clearance of [^64^Cu]Cu-F7D was rapid with the blood peaking at 1 h post-injection, 1.867% ± 0.067, and was four-fold lower than the tumor uptake at 24 h with an average uptake of 0.305% ± 0.048 (Figure 4C,D). *Ex vivo* analysis at 24 h mirrored the ROI analysis demonstrating that [^64^Cu]Cu-F7D exhibited higher uptake in FAP-positive tumors compared to negative controls, with minimal accumulation in non-target tissues aside from expected clearance organs such as the liver and kidneys (Figure S4).

**Figure 4.**
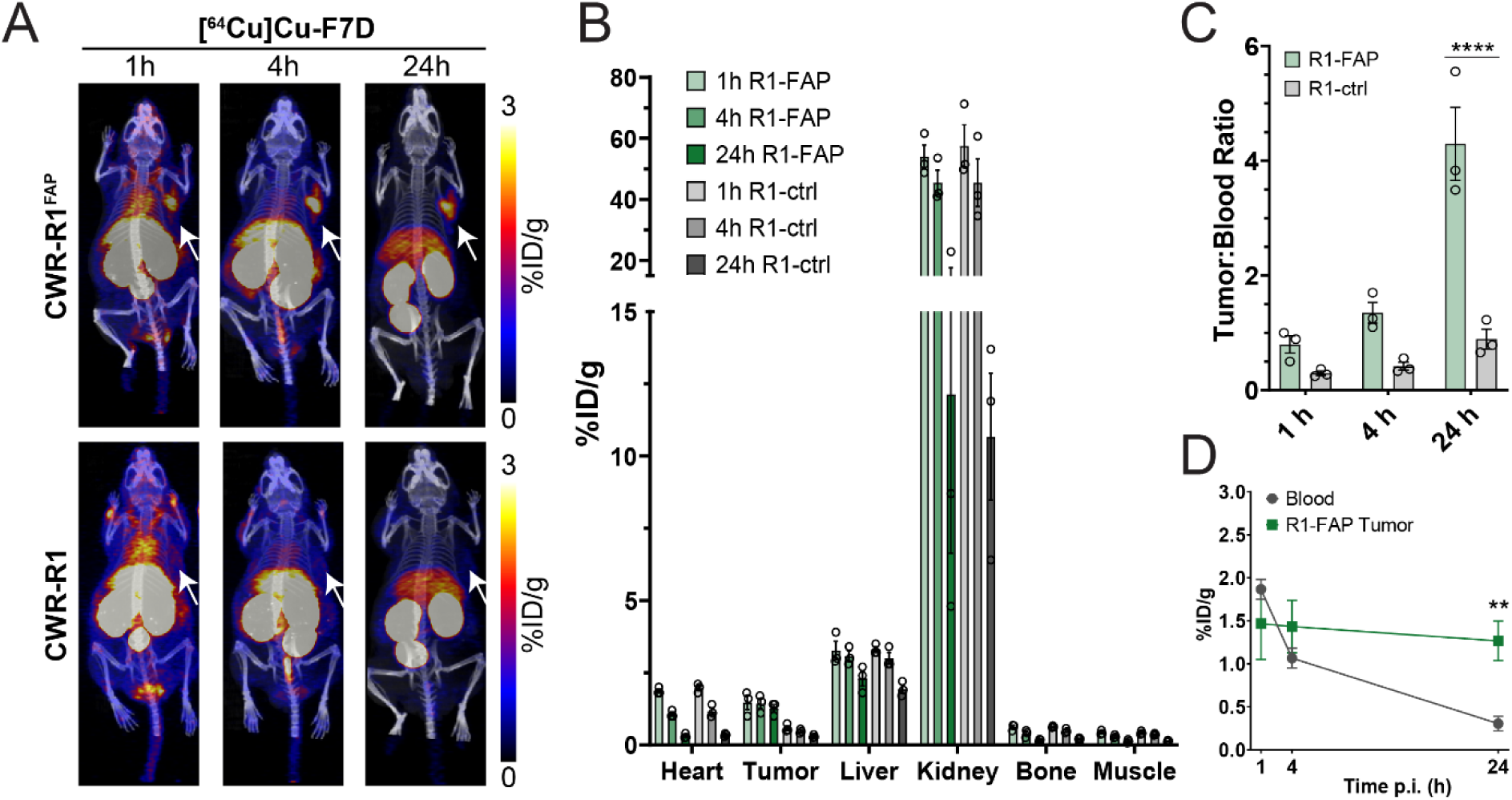
PET imaging and quantitative biodistribution of F7D shows specific and rapid uptake in CWR-R1^FAP^ xenografts. (A) MIPs from µPET/CT scans using [^64^Cu]Cu-F7D in CWR-R1^FAP^ and CWR-R1 xenografts at 1 h, 4 h, and 24 h post-injection. Tumors are indicated by a white arrow. (B) *In vivo* biodistribution among the specified organs at the designated timepoints in mice bearing CWR-R1^FAP^ or CWR-R1 xenografts for [^64^Cu]Cu-F7D. (C) Tumor-to-blood ratio highlighting the clearance and tumor uptake pattern over 24 h. (D) Time-dependent uptake of [^64^Cu]Cu-F7D in blood and CWR-R1^FAP^ tumors, demonstrating sustained tumor retention and clearance from blood over 24 h. *P values*, * *P* ≤ 0.05 ** *P* ≤ 0.01, *** *P* ≤ 0.001, **** *P* ≤ 0.0001 using ordinary two-way ANOVA with Šidák multiple comparisons test.

Compared to F7 and F7D, the largest construct radiolabeled with ^64^Cu ([^64^Cu]Cu-F7-Fc) demonstrated the highest uptake in FAP-positive tumors even at 4 h post-injection (Figure 5A).

**Figure 5.**
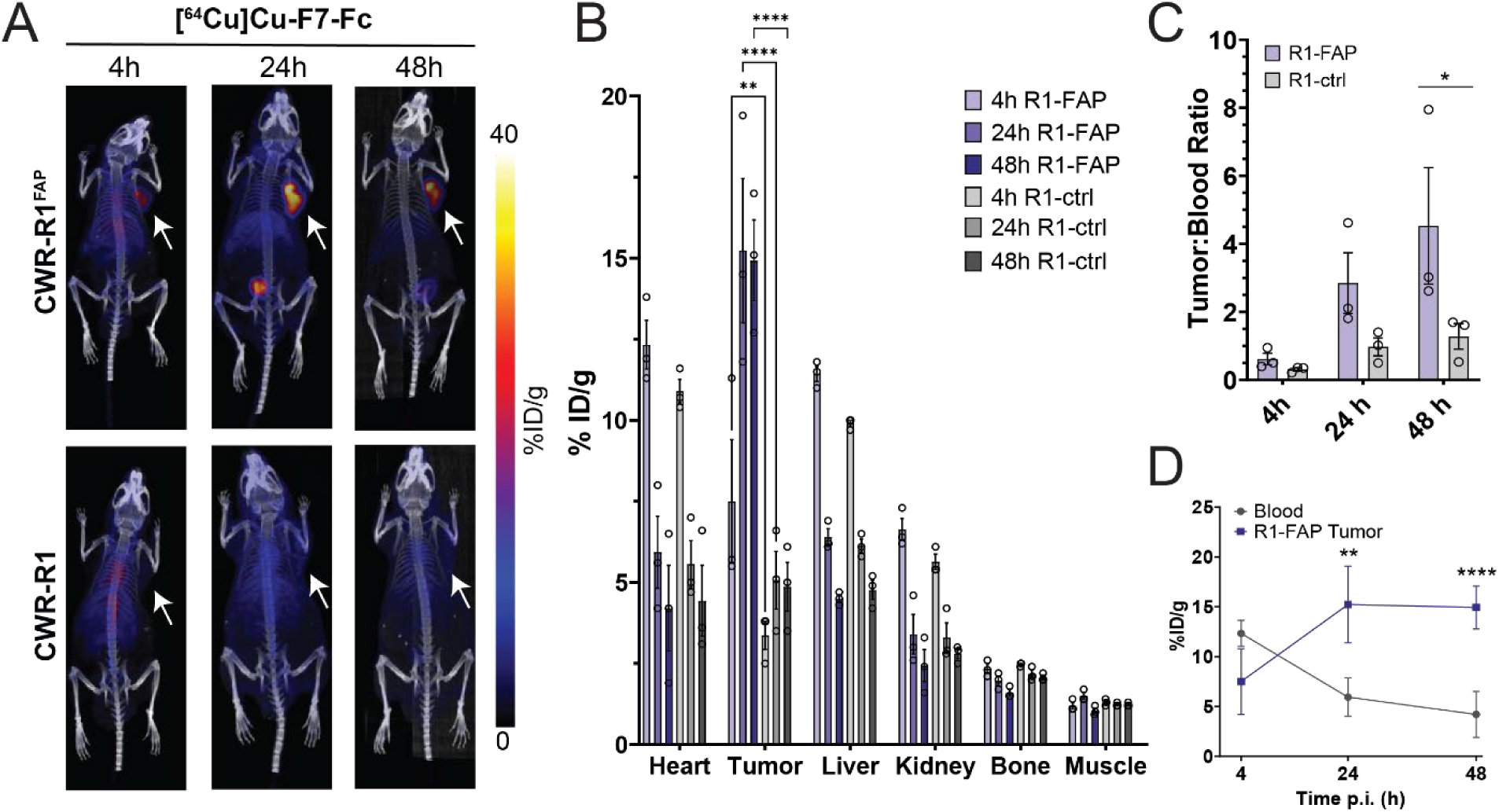
PET imaging and quantitative biodistribution of F7-Fc showing enhanced uptake and retention in CWR-R1^FAP^ xenografts. (A) MIPs from PET/CT scans using [^64^Cu]Cu-F7-Fc in CWR-R1^FAP^ and CWR-R1 xenografts at 1 h, 4 h, and 24 h post-injection. Tumors are indicated by a white arrow. (B) *In vivo* biodistribution among the specified organs at the designated timepoints in mice bearing CWR-R1^FAP^ or CWR-R1 xenografts for [^64^Cu]Cu-F7-Fc. (C) Tumor-to-blood ratio highlighting the clearance and tumor uptake pattern over 24 h. (D) Time-dependent uptake of [^64^Cu]Cu-F7-Fc in blood and CWR-R1^FAP^ tumors, demonstrating sustained tumor retention and clearance from blood over 24 h. *P values*, * *P* ≤ 0.05 ** *P* ≤ 0.01, *** *P* ≤ 0.001, **** *P* ≤ 0.0001 using ordinary two-way ANOVA with Šidák multiple comparisons test.

At 4 h post-injection, tumor uptake in the FAP-positive xenograft was 7.5% ± 1.901. At the 24 h and 48 h timepoints, retention and accumulation of [^64^Cu]Cu-F7-Fc was observed with tumor uptake of 15.23% ± 2.24 and 14.93% ± 1.24 respectively by ROI analysis (Figure 5B). Blood clearance was also apparent with values of 5.93% ± 1.10 and 4.20% ± 1.32 at 24 h and 48 h (Figure 5C,D). *Ex vivo* analysis at 48 h again mirrored ROI analysis further demonstrating [^64^Cu]Cu-F7-Fc specificity and retention in FAP-positive tumors (Figure S5). The most significant difference between [^64^Cu]Cu-F7D and [^64^Cu]Cu-F7-Fc can be seen in the uptake of the liver and kidney. [^64^Cu]Cu-F7D had high uptake in the kidney with 45.53% ± 4.04 at 4 h and 12.13% ± 5.50 at 24 h, whereas the kidney uptake with [^64^Cu]Cu-F7-Fc peaked at 6.63% ± 0.34 at 4 h and was 3.40% ± 0.61 at 24 h. In contrast the liver uptake for [^64^Cu]Cu-F7D was 3.06% ± 0.18 at 4 h and 2.30% ± 0.27 at 24 h. Given its larger size and presence of an Fc domain, the liver uptake of [^64^Cu]Cu-F7-Fc was greater - 11.43% ± 0.233 at 4 h and 6.40% ± 0.25 at 24 h.

To further evaluate the *in vivo* behavior of the Fc-fusion construct, [^89^Zr]Zr-F7-Fc was administered to mice bearing CWR-R1^FAP^ xenografts, and longitudinal PET imaging was performed from 4 h out to 144 h. MIP images revealed progressive accumulation of [^89^Zr]Zr-F7-Fc in FAP-positive tumors with sustained retention over the entire imaging period (Figure 6A).

**Figure 6.**
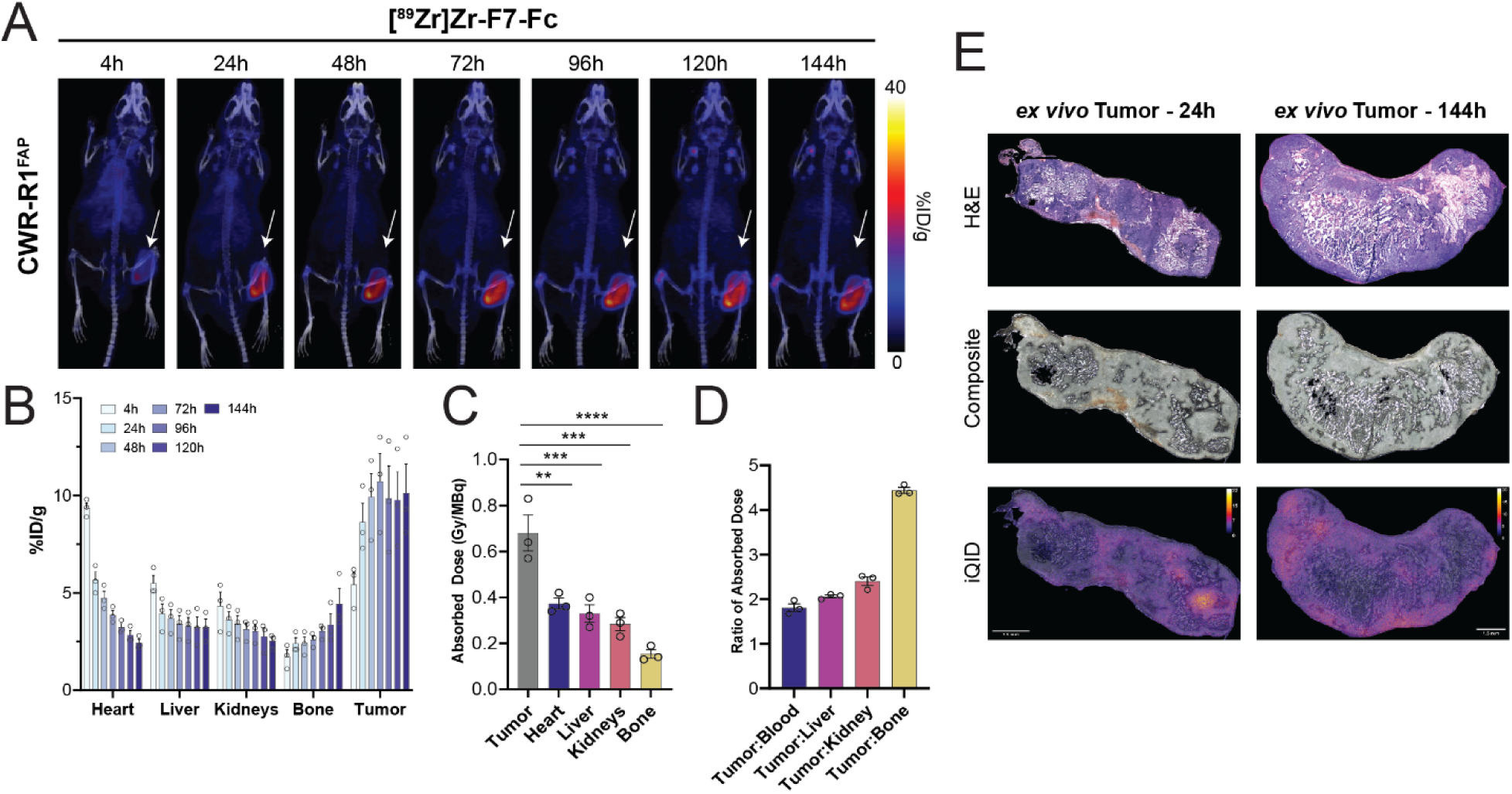
F7-Fc demonstrates prolonged retention in FAP-positive xenografts at 144 h. (A) MIPs from PET/CT scans using [^89^Zr]Zr-F7-Fc in CWR-R1^FAP^ xenografts from 4 h to 144 h post-injection. Tumors are indicated by a white arrow. (B) *In vivo* biodistribution of [^89^Zr]Zr-F7-Fc by ROI analysis of indicated organs at various timepoints in mice bearing CWR-R1^FAP^xenografts for. (C) Estimated absorbed radiation dose (Gy/MBq) for tumor and major organs, showing significantly higher dose delivered to tumor compared to normal tissues. (D) Ratios of absorbed dose between tumor and normal tissues (blood, liver, kidney, bone), demonstrating strong tumor selectivity. (E) *Ex vivo* analysis of tumors at 24 h and 144 h post-injection with H&E staining (top), composite autoradiography (middle), and iQID (bottom), confirming persistent radiotracer localization within tumor tissue over time. *P values*, \**P* ≤ 0.05 ** *P* ≤ 0.01, *** *P* ≤ 0.001, **** *P* ≤ 0.0001 using an ordinary one-way ANOVA with Dunnett’s multiple comparisons test.

Tumor signal became clearly detectable by 24 h, with an uptake of 8.63% ± 1.67, and reaching a peak of 10.73% ± 2.47 at 72 h by ROI analysis. Uptake remained robust at 144 h, at 10.13% ± 2.55, consistent with the prolonged circulation and retention of the Fc-fusion construct (Figure 6B). Absorbed dose calculations confirmed that tumors received significantly higher radiation doses compared to normal tissues (Figure 6C), with tumor-to-organ dose ratios exceeding 4:1 for bone and greater than 2:1 for liver and kidney (Figure 6D). *Ex vivo* analysis confirmed tumor-specific retention of [^89^Zr]Zr-F7-Fc at both 24 h and 144 h timepoints. At 24 h, hematoxylin and eosin (H&E) staining revealed preserved tumor architecture, while composite autoradiography overlays revealed tracer distribution throughout the tissue. iQID analysis confirmed high signal intensity across the tumor, indicating tracer retention at this early timepoint. At 144 h, H&E staining patterns remained largely unchanged, however, composite and iQID images reveal reduced tracer intensity and a heterogeneous distribution, consistent with clearance of the radiotracer and tumor necrosis (Figure 6E). These findings align with *in vivo* imaging data, supporting sustained tumor uptake at prolonged timepoints.

## 4. DISCUSSION

FAP is a well-established TME marker found in a myriad of cancers^31,32^. The biological role of FAP is being uncovered with studies suggesting it may promote immunosuppression, tumor growth, proliferation, and metastasis.^11,33^ Targeting an antigen in the TME may prove critical for the development of diagnostics and therapeutics across multiple solid cancer types.^22^ Thus far, a large effort has been focused on the development of theranostics using small-molecule FAPIs that target the active site of the protease.^22,34^ While similar approaches using small molecules have been successful for other antigens, namely the metalloprotease prostate-specific membrane antigen (PSMA), there are potential pitfalls for the development of these agents for FAP.^35^ The limited molecular interactions FAPIs can make with FAP dictates their affinity as does the choice of warhead on the proline mimetic.^34,36^ As a result, FAPIs exhibit a rapid washout from the tumor which may limit their potential efficacy. FAPIs also require a formed active site with the catalytic triad intact to function.^37^ Endogenous protease inhibitors for FAP may exist and dimerization of the protein is required for activity.^37^ FAP is also known to form dimers with dipeptidyl peptidase-4 (DPPIV) on the cell surface.^33^ For these reasons, we aimed to develop a protein-based approach utilizing small VHHs. These proteins can be developed to recognize sites distant or adjacent to the active site and can mimic the kinetics of small molecules, yet with greater uptake and retention.

In this study, we identified a novel VHH for FAP that allowed us to engineer multiple targeting vectors with different pharmacokinetic and antigen engagement properties. The small monomeric VHH F7 was able to compete with the clinically relevant FAPI-46. At 1 h post-injection, the tumor uptake of F7 was nearly two-fold that of FAPI-46 and this uptake was retained out to the last timepoint at 4 h. The retention of F7 by the tumor allowed for a reduction in the blood pool signal and the renal uptake to decline. This profile of quick tumor uptake, retention, and clearance underscores how monomeric VHHs are similar to small molecules. The targeting potential of F7 became more apparent as the molecular weight of the constructs increased. While the molecular weight of F7D was only around twice that of F7 VHH, the extra avidity of two VHHs tethered together saw the K_D_ improve more than 300-fold to 29pM. This allowed F7D to rapidly localize to the FAP-positive xenograft and be retained likely due to the extremely slow K_d_. The tumor-to-blood ratio of the dimer improved significantly between 1 h and 4 h with further improvement at 24 h. The Fc-fusion construct, F7-Fc, demonstrated markedly superior tumor uptake and retention compared to its lower molecular weight counterparts, F7 and F7D, highlighting the influence of molecular size and Fc incorporation on pharmacokinetics and biodistribution. In longitudinal PET imaging with ^89^Zr labeling, F7-Fc exhibited detectable tumor uptake as early as 4 h post-injection and remained in the tumor at the final timepoint of 144 h post-injection. Across both ^64^Cu and ^89^Zr studies, the construct consistently maintained high tumor retention while showing progressive clearance from blood and catabolic organs such as the liver and kidneys. This favorable biodistribution profile suggests that Fc-VHH fusions may reduce off-target toxicity commonly associated with traditional radioimmunotherapies, while retaining strong potential as standalone imaging agents.

## 5. CONCLUSIONS

In a first of its kind study, we investigated what effects valency and molecular weight had on the ability of a single-domain VHH to target FAP. As valency changed from a single monomer to bivalent constructs, an increase in affinity was observed with concomitant tumor retention *in vivo* largely driven by a decreased dissociation rate. The highest molecular weight construct, F7-Fc, had high tumor uptake retention which allowed for longitudinal imaging out to 144 h. Future studies need to be performed to see how these properties affect therapeutic efficacy *in vivo*.

## ASSOCIATED CONTENT

### Supporting Information

The Supporting Information is available free of charge. Supplemental data consists of Supplemental Methods (file type: DOCX), 6 figures (file type: DOCX), and 2 tables (file type: DOCX) Supplemental Methods provide study-relevant details on phage display biopanning, ELISA, protein purification, bio-layer interferometry, and flow cytometry. Figures present data on identification of anti-hFAP NbC VHH from a naïve camelid antibody phage display library (Figure S1); identification of anti-hFAP NbC VHH from an affinity-matured camelid antibody phage display library (Figure S2); *ex vivo* biodistribution of [^64^Cu]Cu-F7D among blood and specified organs and tissues (Figure S3); and *ex vivo* biodistribution of [^64^Cu]Cu-F7-Fc among blood and specified organs and tissues (Figure S4). Tables present data on unique anti-hFAP VHH clones isolated from a naïve camelid antibody phage display library (Table S1) and unique anti-hFAP VHH clones isolated from an affinity-maturated camelid antibody phage display library (Table S2).

## AUTHOR INFORMATION

### Notes

The authors declare the following competing financial interest(s): Aaron M. LeBeau and Joseph P. Gallant are listed as inventors on a provisional patent regarding this research (US Provisional Application 63/531,45).

## ACKNOWLEDGMENTS

Funding: This work was supported by NIH/NCI R01 CA237272 (A.M.L.), NIH/NCI R01 CA233562 (A.M.L.), a Prostate Cancer Foundation Challenge Award (A.M.L.), a Prostate Cancer Foundation Young Investigator Award (A.M.L.), and Andy North and Friends (A.M.L.). We thank the staff at the Small Animal Imaging and Radiotherapy Facility at the University of Wisconsin School of Medicine and Public Health for their assistance. The authors would also like to thank James P. Zacny for manuscript formatting and editing assistance.

## Notes

### Competing Interest Statement

The authors have declared no competing interest.

