## Supplemental Information for "Tuning the Structural Properties of a Single-Domain Antibody Scaffold for Improved Fibroblast Activation Protein Targeting"

### Supplemental Methods

**Phage display biopanning.** In order to identify a high affinity lead, fibroblast activation protein (FAP)-targeting variable-heavy-heavy domain (VHH), a phage biopanning campaign was performed. Starting from a previously identified lead, affinity maturation was conducted using a controlled mutational scheme (GeneART, ThemoFisher) generating a library size of ~$10^8$ with an average amino acid mutational rate of 1 in the CDR1 region and 4 in the CDR3 region. Variants were packaged into a phagemid (PADL22c, Antibody Design Labs), rescued with M13K07 helper phage, and subjected to two rounds of biopanning following established protocols [1].

**ELISA.** FAP-targeted VHHs were produced by SS320 *E. coli* from a total of 864 individual clones using 5mM IPTG induction in a 96-well growth plate. The supernatant following induction and overnight incubation at 30°C with 250 rpm were screened for binding to hFAP by ELISA. MaxiSorp Nunc plates were coated with 5ug/mL of streptavidin overnight at 4°C. The wells were washed twice with PBS before the wells were blocked with 370 µl of 2% BSA for 1 h at room temperature on a small radius shaker with 300 rpm. Unless noted, all additional incubations are at room temperature with 300 rpm shaking. A subsequent wash was performed three times with PBST (0.005% Tween20). The recombinant FAP (ACRO FAP-H82Q6) was added to the wells at 1µg/mL in PBST with 1% BSA and allowed to incubate for 1 h. Wells were washed 3 times again to remove any unbound FAP before the supernatants of the VHH clones was added to the wells diluted in PBST with 1% BSA and allowed to incubate for 1 h. Wells were washed again with PBST 3 times before an anti-HA HRP conjugated antibody (Sigma Aldrich 12-013-819-01) was added in a 1:1000 dilution in PBST with 1% BSA and incubated for 1 h. For the final visualization of the VHH binding, the wells were washed three times with PBST before 50µl of Turbo TMB reagent (Pierce) was added to each well. The peroxidase reaction was allowed to occur for no longer than 5 min and was stopped with 10µl of 1M $H_2SO_4$. The absorbance at 450 nm was read using a microplate reader. Positive clones underwent a second ELISA using a dilution series of induced supernatant and the top hits were sequenced to identify unique clones.

**Protein purification**. The VHH phagemid was cloned into a modified pet22b expression vector to include a TEV protease cleavable site to remove the 6x His purification tag. The plasmid containing was cloned into Shuffle T7 *E. coli* cells and grown in TB media. For purification 1 L of F7 containing *E. coli* was grown to $OD_{600}$ of 0.8 at 37°C with 250 rpm before being inoculated with 10 mM IPTG and once induced was grown at 16°C with 250 rpm overnight. The following day the F7 VHH was purified using a periplasmic prep method. Briefly the cells were spun down at 7500xRCF for 15 min and supernatant was removed. The cell pellet was resuspended in 30 mL of TES buffer and slowly stirred for 30 min at 4°C. The cells were spun again at 7500xRCF for 30 min and the pellet was resuspended in 30 mL of Tris Buffer and stirred slowly at 4°C for 30 min. The cells went through a final spin at 7500xRCF for 30 min. The supernatant containing the VHH was collected and captured via a 5 mL HiTrap HP His column (Cytiva) before an imidazole purification was run on an AKTA HPLC. The purified VHH was collected, and TEV protease was used to cleave the affinity tags, and the sample was rerun over the His column, with flowthrough collected and the final protein sample was dialyzed into PBS. The F7 dimer was produced in a similar fashion with the change that the modified plasmid contained a 5x GS4 linker with the F7 VHH cloned onto either flanking side of the linker and a N-terminus TEV cleavable SUMO tag with a 6x His tag. For the F7 dimer, once purified via a 5 mL HiTrap HP His column, the protein was combined with a TEV protease to remove the 6xHis-SUMO tag and VHH was purified again collecting the flowthrough of the 5 mL HiTrap HP His column followed by size exclusion chromatography (HiLoad Superdex 75 10/300 (Cytiva)).

Vectors for mammalian expression of F7-Fc were generated by Genscript. Codon-optimized sequences encoding F7 were synthesized *de novo* and subcloned into TGEX-SCblue (Antibody Design Labs) in frame with the PelBK and huG1-Fc sequences, using standard molecular biology methods. Constructs were transformed into competent NEB 5-alpha cells (New England Biolabs) and plasmid DNA was purified using PureLink HiPure plasmid maxiprep kits (Thermo Fisher) according to the vendor's recommended protocol. For the F7-Fc protein production, ExpiCHO-S cells were maintained in ExpiCHO Expression Medium at 37°C and 8% CO2, shaking at 120 rpm on a 25 mm orbital diameter, in vented Erlenmeyer flasks. Cells were cultured to a density of $6x10^6$ cells/ml with a viability >95% and were transfected with F7-TGEX-SCblue. Transfections were performed using an Expifectamine CHO Transfection Kit (Thermo Fisher) according to the vendor's recommended protocol, using 1 µg of plasmid DNA and 3.2 µl of Expifectamine CHO Reagent per ml of suspension culture. Cells were returned to 37°C and 8% CO2 incubator overnight, shaking at 120 rpm. The following day, transfected cells were supplemented with 6 µl of ExpiCHO Enhancer and 240 µl of ExpiCHO Feed per ml of suspension culture, and incubator conditions were changed to 32°C and 5% $CO_2$. Twelve days post-transfection, suspension cultures were harvested and centrifuged at 2000xRCF for 10 min at 4°C. The protein-containing supernatant was collected and further clarified by centrifuging at 20,000xRCF for 30 min at 4°C and passed through a 0.22 µm sterile filter. To purify the F7-Fc antibody it was captured using protein A and was further purified by size exclusion chromatography. A HiTrap Protein A HP column (Cytiva) was equilibrated with 5 column volumes (CVs) of PBS. Clarified ExpiCHO-S supernatant was supplemented with 300mM NaCl, and pH was adjusted to 6.8 and flowed through a protein A column at 1CV/min. Unbound protein was washed from the column using 10 CVs of PBS. F7-Fc was eluted in 2.5 CVs of 100mM glycine pH 3.0, followed by 2.5 CVs of PBS. Eluate was immediately neutralized using 2M Tris HCl pH 8.6. Eluate was concentrated and buffer exchanged into PBS using an Amicon Ultra Centrifugal Filter with a 30kDa MWCO membrane (Millipore). Size exclusion chromatography was performed on an ÄKTApure fast protein liquid chromatography system using a HiLoad 16/600 Superdex 200 PG (Cytiva) column equilibrated with PBS. Samples were loaded and fractionated using a mobile phase of PBS at 0.5 ml/min, and chromatograms were obtained by monitoring UV absorbance at 280 nm. Eluted protein fractions contributing to a single peak of UV absorbance corresponding to the theoretical molecular mass of F7-Fc were collected and pooled. Eluates were diluted to 1 mg/ml in PBS, aliquoted, and flash-frozen for storage.

**Bio-layer interferometry.** Bio-layer interferometry measurements were obtained using a Sartorius Octet R8. BLI experiments followed established protocols.[1,2] Briefly, biotinylated human FAP protein (ACRO FAP-H82Q6) was captured on hydrated SAX (high precision streptavidin) biosensors. An assay buffer of PBS with 1% BSA was used for all steps including dilutions. Hydrated SAX biosensors were equilibrated for 30 s in the assay buffer before the biotinylated FAP was captured for 45 s. A second equilibration step was carried out for 30 s. Serial dilutions of the various VHH constructs were then exposed to the FAP-loaded SAX biosensor for 180 – 240 s followed by a dissociation step in assay buffer for an equivalent amount of time. Two separate controls were run to evaluate non-specific binding. One control included a FAP-loaded SAX biosensor exposed to assay buffer in both association and dissociation steps. A second control included a SAX biosensor left unloaded with FAP and exposed to the highest concentration of the VHH constructs. The latter control well was subtracted from the data before modeling and the dissociation constant $K_D$ was calculated. Graphs were drawn using the corrected data in GraphPad Prism.

**Flow Cytometry.** CWR-R1 and CWR-R1$^{FAP}$ cell lines were harvested and resuspended in flow cytometry staining buffer (eBioscience, Invitrogen). Cells were incubated with Alexa Fluor 647–labeled VHH constructs at concentrations ranging from 250 - 0.24 nM for 1 hour on ice. Following incubation, cells were centrifuged at 400xRCF for 4 min and washed three times with staining buffer. The cell pellets for each sample were then resuspended in 300 µL of staining buffer and analyzed using an Attune Flow Cytometer (Thermo Fisher Scientific). Fluorescence intensity along with histogram generation was performed using FlowJo software (Tree Star, Inc.).

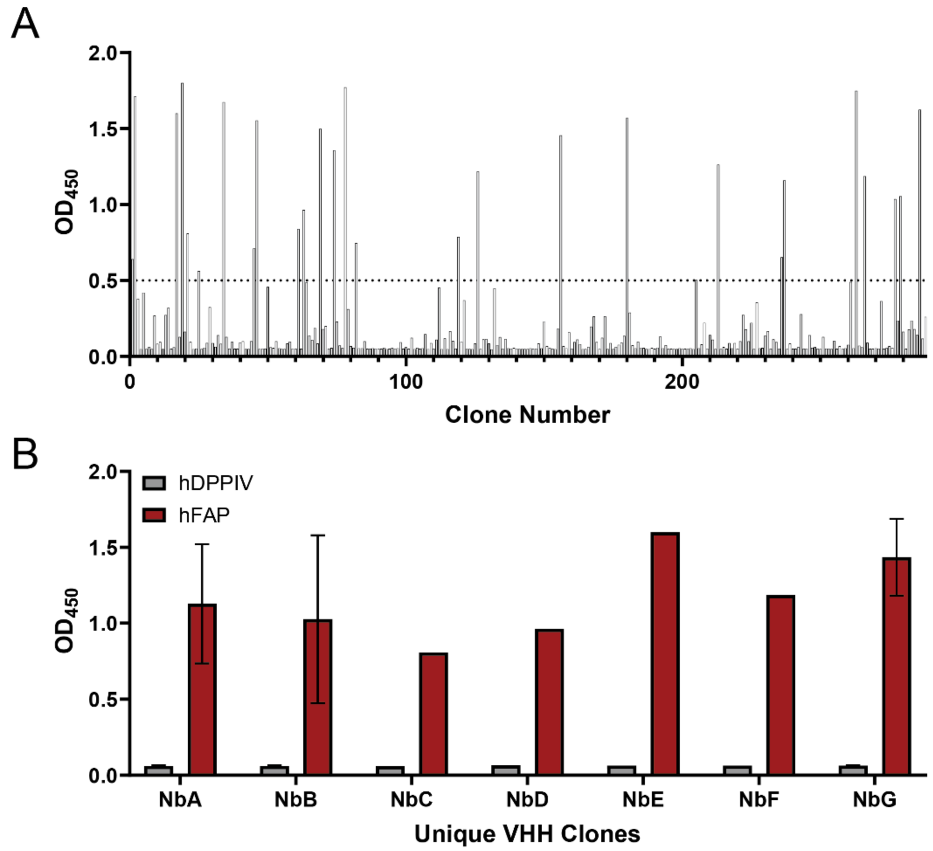


**Figure S1.** Identification of anti-hFAP NbC VHH from a naïve camelid antibody phage display library. (A) A total of 288 unpurified VHH clones from culture supernatants were screened on hFAP-coated plates. Clones with $OD_{450}$ ≥ 0.5 were considered strong binders. (B) Unique VHH clones demonstrating hFAP specificity compared to closely related protein, hDPPIV, by ELISA.

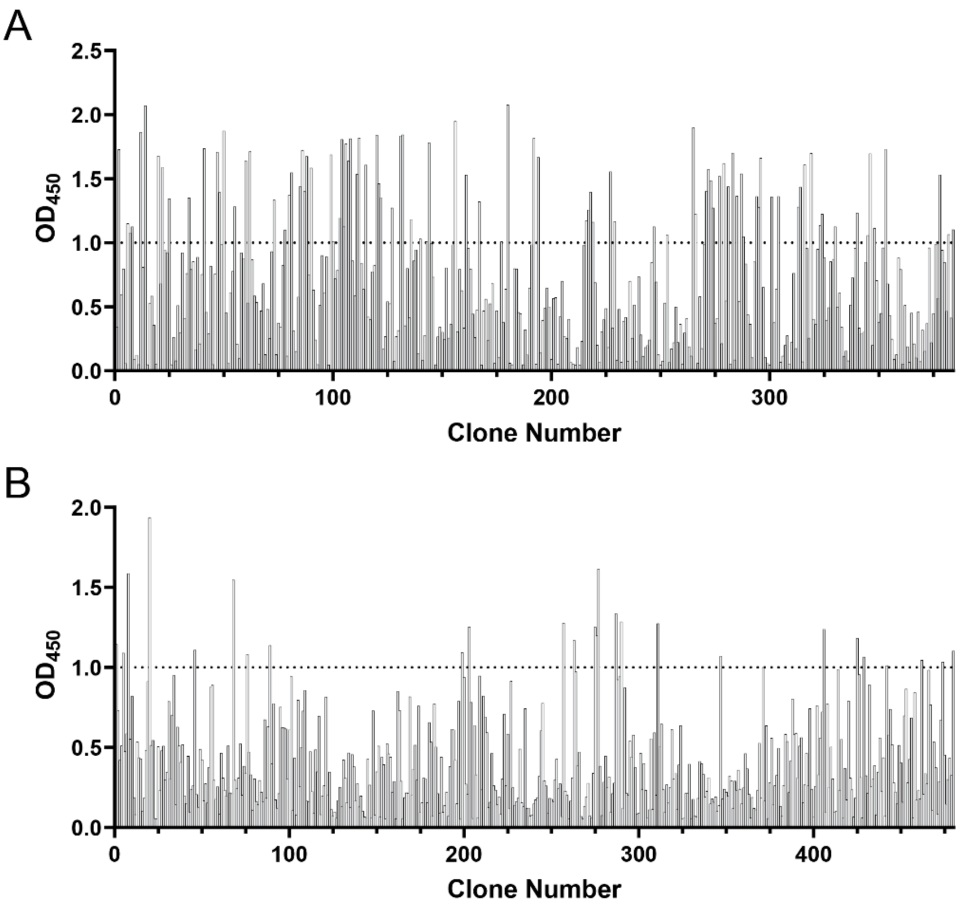


**Figure S2.** Identification of anti-hFAP NbC VHH from an affinity-matured camelid antibody phage display library. Screening of unpurified VHH clones by ELISA on hFAP-coated plates for biopanning rounds 5 (A) and 6 (B) for a total of 864 clones. Clones with $OD_{450}$ ≥ 1.0 were considered strong binders.

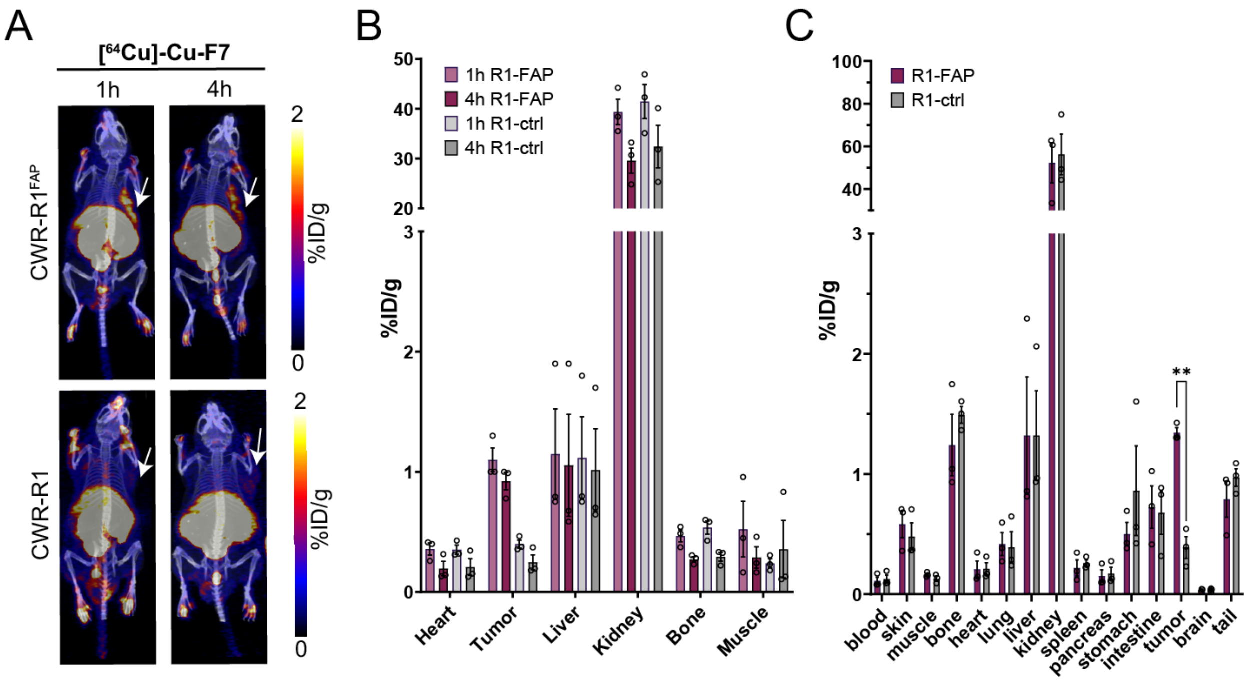


**Figure S3.** Whole-body PET/CT imaging, quantitative PET analysis, and biodistribution of F7. (A) maximum intensity projections (MIPs) from PET/CT scans using [$^{64}$Cu]Cu-F7 in CWR-R1$^{FAP}$ and CWR-R1 xenografts at 1 h and 4 h post-injection. Tumor indicated by white arrow. (B) *In vivo* biodistribution among the indicated organs at the indicated timepoints in mice bearing CWR-R1$^{FAP}$ or CWR-R1 xenografts for [$^{64}$Cu]Cu-F7. (C) *Ex vivo* biodistribution of [$^{64}$Cu]Cu-F7 among blood and specified organs and tissues. *P values*, * $P \leq 0.05$, ** $P \leq 0.01$ using unpaired T test with Welch correction.

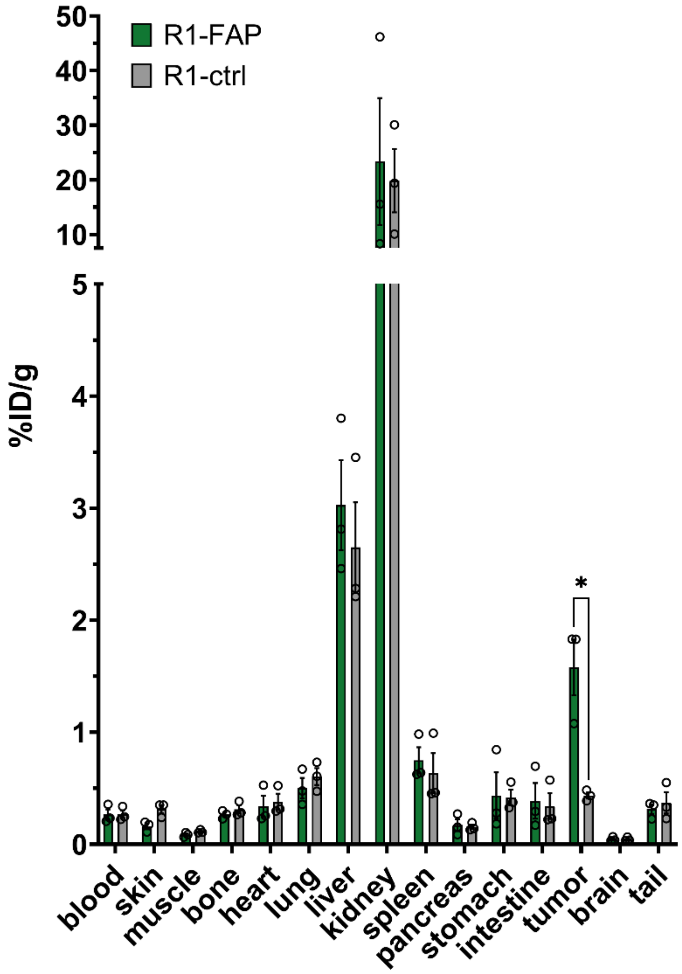


**Figure S4.** *Ex vivo* biodistribution of [$^{64}$Cu]Cu-F7D among blood and specified organs and tissues. *P values*, * *P* ≤ 0.05, ** *P* ≤ 0.01 using unpaired T test with Welch correction.

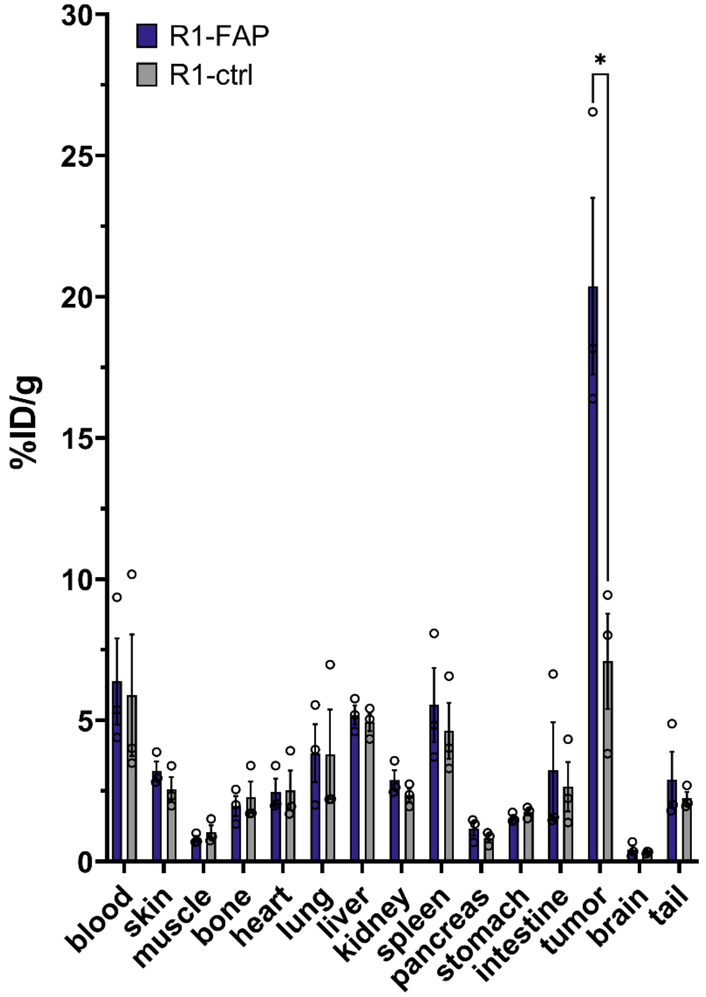


**Figure S5.** *Ex vivo* biodistribution of [$^{64}$Cu]Cu-F7-Fc among blood and specified organs and tissues. *P values*, *P ≤ 0.05, ** *P* ≤ 0.01 using unpaired T test with Welch correction.

| Clone ID | Sequence |
|---|---|
| NbA | QVQLVESGGGLVQAGGSLRLSCTASGSPFRLNVMGWYRQAPGKQRELVADATSGGSTNYADIVKGRFTISRDNAKNRVYLQMDSLKPEDTAVYYCAASSTLDPRRYYSGDYSPDEYDFWGQGTQVTVS |
| NbB | QVQLVESGGGLVQAGGSLRLSCTASGSPFRLNVMGWYRQAPGKQRELVADATSGGSTNYADIVKGRFTISRDNAKNRVYLQMDSLKPEDTAVYYCAASSTLDPRRYYSGDYSPDEYDFWGQGTLVTVS |
| NbC | QVQLVESGGGLVQPGGSLRLSCAASGFTFNNYWVYWARQAPGKGLEWVSSMNIADGTTYYAASVKGRFTISSDNAKNTVYLQMNSLKSEDTAVYYCAKGVGGPVGTGGHYLGKGTLVTVS |
| NbD | QVQLVESGGGLVQAGGSLRLSCAISGVTSSIGTTAWHRQAPGKQRELVALLVNDGRTVYNEAVKGRFTISRDNAKGTVDLQMNVLEAEDTAVYYCYGDTYGTPLVDHKKYWGKGTLVTVS |
| NbE | QVQLVESGGGLVQPGGSLRLSCAVSGFTFSTYGMSWVRQAPGKGLEWVSYINSGGSYTRYADSVKGRFTISRDNANNMLYLQMNSLEPEDTAVYYCSNRDSWGQGTLVTVS |
| NbF | EVQLVESGGGSVRTGGSLTLSCIISGFTFNDYSMFWVRQAPGKGLEWVASISAGGATTLYRDSVKGRFTISSDGAKKTVTLQMNDLKPEDTALYYCAADPNPAVARGSPRRVPWLKGQGTQVTVS |
| NbG | QVQLVESGGGTVQPGGSLRVSCAASGFTFSNYAMSWVRQAPGKGPEWVSGISDGGRTYYRYSVKGRFTISRDNAKNTLYLQMNNLKVEDTALYYCARAPADPTYGMDYWGKGTLVTVS |

**Table S1.** Unique anti-hFAP VHH clones isolated from a naïve camelid antibody phage display library.

| Clone ID | Sequence |
|---|---|
| R2P2 F7 | QVQLVESGGGLVQPGGSLRLSCAASGFTFNNYWVYWARQAPGKGLEWVSSMNIADGTTYYAASVKGRFTISSDNAKNTVYLQMNSLKSEDTAVYYCAKGVVGPVGNGGLYLGKGTLVTVS |
| R2P2 G6 | QVQLVESGGGLVQPGGSLRLSCAASGFTFNNYWVYWARQAPGKGLEWVSSMNMADGTTYYAASVKGRFTISSDNAKNTVYLQMNSLKSEDTAVYYCAKGDGGVVGSGGHYLGKGTLVTVS |
| R2P2 H1 | QVQLVESGGGLVQPGGSLRLSCAASGFTFNNYWVYWARQAPGKGLEWVSSMNIADGTTYYAASVKGRFTISSDNAKNTVYLQMNSLKSEDTAVYYCANGVGGIVGTGGHYLGKGTLVTVS |
| R2P3 F11 | QVQLVESGGGLVQPGGSLRLSCAASGFTFNNYWVYWARQAPGKGLEWVSSMNIADGTTYYAASVKGRFTISSDNAKNTVYLQMNSLKSEDTAVYYCAKGVVGPVGNGGLYLGKGTLVTVS |
| R2P3 H1 | QVQLVESGGGLVQPGGSLRLSCAASGFTFNNYWVYWARQAPGKGLEWVSSMNIADGTTYYAASVKGRFTISSDNAKNTVYLQMNSLKSEDTAVYYCAKGVGGPLGSGGHYLGKGTLVTVS |

**Table S2.** Unique anti-hFAP VHH clones isolated from an affinity-maturated camelid antibody phage display library.
